# A Parallel Ratchet-Stroke Mechanism Leads to an Optimum Force for Molecular Motor Function

**DOI:** 10.1101/2020.06.29.177964

**Authors:** U.L. Mallimadugula, E.A. Galburt

## Abstract

Molecular motors convert chemical potential energy into mechanical work and perform a great number of critical biological functions. Examples include the polymerization and manipulation of nucleic acids, the generation of cellular motility and contractility, the formation and maintenance of cell shape, and the transport of materials within cells. The mechanisms underlying these molecular machines are routinely divided into two categories: Brownian ratchet and power stroke. While a ratchet uses chemical energy to bias thermally activated motion, a stroke depends on a direct coupling between chemical events and motion. However, the multi-dimensional nature of protein energy landscapes allows for the possibility of multiple reaction paths connecting two states. Here, we investigate the properties of a hypothetical molecular motor able to utilize parallel ratchet and stroke translocation mechanisms. We explore motor velocity and force-dependence as a function of the energy landscape of each path and reveal the potential for such a mechanism to result in an optimum force for motor function. We explore how the presence of this optimum depends on the rates of the individual paths and show that the distribution of stepping times characterized by the randomness parameter may be used to test for parallel path mechanisms. Lastly, we caution that experimental data consisting solely of measurements of velocity as a function of ATP concentration and force cannot be used to eliminate the possibility of such a parallel path mechanism.

**SIGNIFICANCE:** Molecular motors perform various mechanical functions in cells allowing them to move, replicate and perform various housekeeping functions required for life. Biophysical studies often aim to determine the molecular mechanism by which these motors convert chemical energy to mechanical work by fitting experimental data with kinetic models that fall into one of two classes: Brownian ratchets or power strokes. However, nothing *a priori* requires that a motor function via a single mechanism. Here, we consider a theoretical construct where a motor has access to both class of mechanism in parallel. Combining stochastic simulations and analytical solutions we describe unique signatures of such a mechanism that could be observed experimentally. We also show that absence of these signatures does not formally eliminate the existence of such a parallel mechanism. These findings expand our theoretical understanding of the potential motor behaviors with which to interpret experimental results.

## INTRODUCTION

Biology is replete with macromolecular enzymes that convert chemical potential energy into mechanical work. These enzymes come in both protein and nucleic acid forms and drive DNA (1), RNA (2–4), and protein polymerization (5) and metabolism (6, 7), muscle contraction (8), cellular transport (9), and flagella rotation (10). In contrast to classical enzymes, which proceed through cycles of chemical catalysis and simply release liberated free energy into the thermal bath, molecular motors have evolved coupled mechanisms that allow for some of the liberated energy to be used in the form of mechanical work (11–14). These mechanisms consist of a series of connected structural intermediates and must account for coupling between chemical (i.e. ATP concentration) and mechanical (i.e. force) variables and are typically divided into two classes: Brownian ratchet or power stroke.

Brownian ratchet mechanisms lead to biased motion by rectifying thermal fluctuations via chemical reactions. For example, the free 1D diffusion of a particle may be rectified via the asymmetry of a flashing ratchet potential (11, 12, 14–17). The asymmetry of the potential results in forward excursions being ratcheted forward, while backwards excursions being ratcheted to the original position. A biological example of a ratchet is perhaps best exemplified by multi-subunit RNA polymerases. These enzymes translocate along a DNA template one base-pair at a time and concomitantly synthesize a complementary RNA transcript. They have been extensively studied by both structural and high resolution single-molecule techniques and the data are consistent with translocation via a Brownian ratchet (3, 18–22). More specifically, during active translocation, the enzyme is found to be in one of two states in the absence of the next nucleotide. In the pre-translocated state, the binding site for the next nucleotide is obscured by the last nucleotide incorporated. In the post-translocated state, the binding site has been vacated via the thermally stimulated movement of the polymerase one base-pair downstream. The model stipulates that in the absence of the next nucleotide a dynamic equilibrium exists between these two states. The incorporation of the next nucleotide into the growing 3’ end essentially converts the post-translocated state into the pre-translocated state and the system is ratcheted forward. Other examples of molecular motors thought to proceed via ratchet mechanisms include the phi29 bacteriophage DNA polymerase (1), the ribosome (5), and some (e.g. passive) helicases (23–25).

In contrast to the ratchet model, power stroke models essentially contain a single kinetic step which directly couples a chemical step to motion (26). This step can be thought of as proceeding diagonally on a 2D energy landscape where one dimension represents distance translocated and another dimension represents progress along the chemical reaction coordinate (27). An example of this behavior can be found in the mechanism of myosin motors translocating along microtubules (28–30). Here the binding of ATP to a free myosin head is followed by ATP hydrolysis. Subsequent actin binding stimulates phosphate release and a concomitant force-generating conformational change or stroke. Other examples of molecular machines thought to be governed by power stroke mechanisms can be found in ATP synthase (31), AAA+ proteases (32) and the viral DNA packaging motor from the bacteriophage phi29 (33).

However, the distinction between ratchet and power stroke function is not always sharp and these definitions may not be as clear as they appear on the surface when applied to real systems. In particular, depending on the magnitude of free energy changes along the pathway of translocation, a ratchet with many small steps may behave very much according to a power stroke (17). In addition, motors like myosin may perform a fraction of their step as a stroke followed by a more ratchet-like mechanism for the subsequent fraction (34).

We recently suggested an analogy between these motor mechanisms and the dual mechanisms of ligand binding: conformational selection and induced fit (35). With ligand binding in the context of a thermodynamic cycle, increasing ligand concentration titrates the equilibrium flux of the system towards the induced fit mechanism (36). Under some circumstances, this effect has a specific consequence that can be experimentally observed. In particular, a biphasic third observed rate of equilibration can be taken as direct evidence of the presence of both conformational selection and induced fit mechanisms working in parallel (37). Here we explore this analogy between ligand binding and motor function further by determining the behavior of a hypothetical motor that is capable of performing via power stroke or Brownian ratchet mechanisms in parallel kinetic pathways (Fig. 1). Surprisingly, under certain conditions, the velocity of such a motor peaks at an optimum force. This unique feature could, in principle, be used as experimental evidence for motor behavior of this kind. We further show that trends in the randomness parameter calculated from single molecule dwell times provides a more sensitive indication of parallel paths. Lastly, we demonstrate that, just as kinetic data cannot rule out the presence of parallel binding paths (37), even force-velocity relationships and single molecule dwell times may not be sufficient to formally eliminate the possibility of a parallel path mechanism.

**Figure 1:**
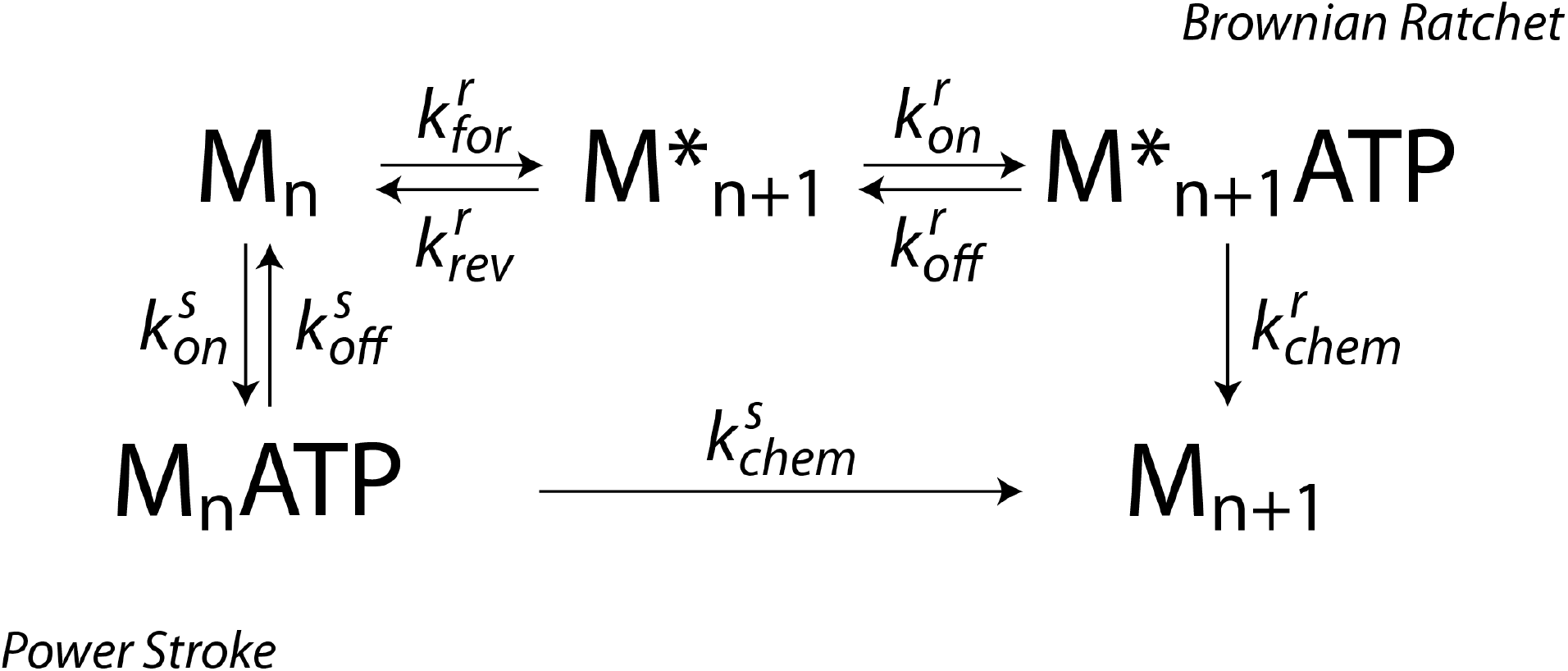
A Parallel Path Molecular Motor Mechanism. The states represent the pre- or post-translocated states of the motor and the ATP-bound state. The rate constants are shown where a superscript “r” refers to the ratchet path and a superscript “s” refers to the stroke path. The chemical steps in both cases include hydrolysis and product release.

## METHODS

Molecular motors typically use a chemical energy source, such as a nucleotide triphosphate molecule or a proton gradient, to drive translocation. We constructed a parallel ratchet-stroke mechanism that uses adenosine triphosphate (ATP) as its energy source and allows for a motor (M) to function via two pathways (Fig. 1). In the ratchet path, M takes a thermally activated and reversible step forward prior to ATP binding and hydrolysis. In the stroke path, M reversibly binds ATP prior to hydrolysis and stepping forward. F is the applied force on M that produces a work (W) given by *W* = *F*Δ*x* where Δ*x* denotes step size of the motor. The effect of this work is incorporated into the force dependent reactions as follows.

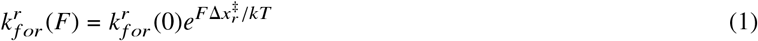

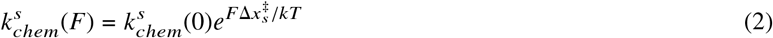

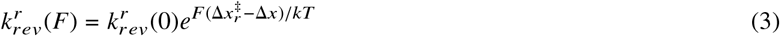

Here, 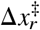 and 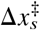 are the distances to the transition state in the forward direction for the ratchet and stroke mechanisms respectively. *k* is Boltzmann’s constant and *T* is temperature. Therefore, the parameters that can be varied in the system include all the basal rate constants shown in Fig.1, applied external force *F*, distance to transition state of the forward translocation reactions of the ratchet 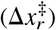 and the stroke 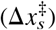 and the concentration of ATP. Simulations, calculations, and analyses were done at *T* = 298*K* using Python 3.6.8 (Anaconda Inc.).

### Stochastic Simulation of Parallel Pathway

To simulate the system, we used the Gillespie algorithm (38, 39) which can be described as follows. For a system with *μ* = (1, 2…8) reactions, the algorithm carries out a stochastic simulation based on the Reaction Probability Distribution function (Eq. 4). This function gives the probability that, given the current state of the system at time t, the next reaction will occur in the interval (*t* + τ, *t* + τ + *d*τ) and will be reaction *μ*.

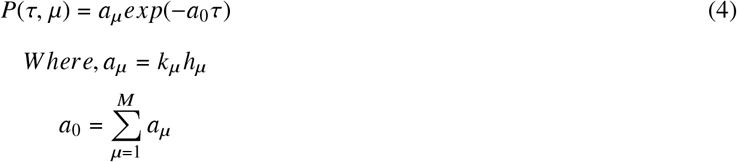

*h_μ_* is the distinct molecular reactant combinations available for reaction *μ* and *k_μ_* is the rate constant for reaction *μ*.

At every iteration the simulation picks a random number from a Poisson distribution of reaction time probabilities. This is used as the waiting time for the next reaction. Additionally, a picks a second random number which is used to determine which of the set of possible reactions occurs. The populations of states are updated and the process is repeated for a sufficient number of iterations to approach a steady-state solution of the process.

### Analytical Steady State Solution

The analytical solution is obtained by solving for the steady state concentrations of each of the species of the system. At steady state, the net flux of each of the species, M, M*, M*.ATP and M.ATP is zero.

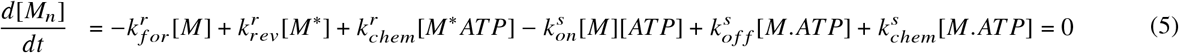

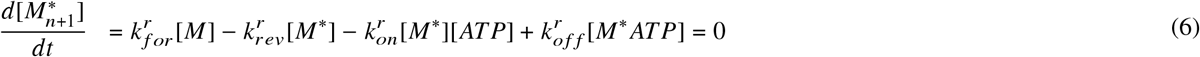

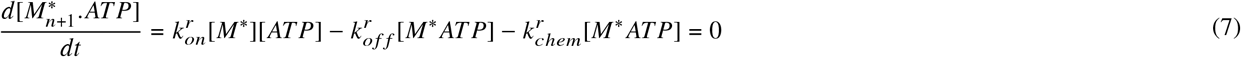

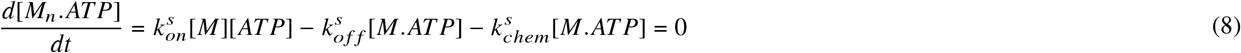

Additionally,

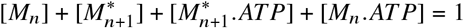

The concentrations obtained by solving the above system of equations are then used to obtain the fractional flux through the ratchet arm (*ϕ*_*r*_) of the mechanism.

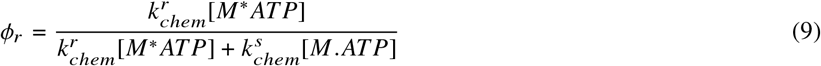

The concentrations in the above equation can be written as combinations of the rate constants and the ATP concentration. This expression is shown in equation 9 in the supplemental information.

*ϕ*_*r*_ and the fractional flux through stroke arm (*ϕ*_*s*_ = 1 − *ϕ*_*r*_) can then be used to obtain an expression for the average motor velocity (⟨*V*⟩):

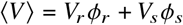

where *V_r_* and *V_s_* are the net rates for the ratchet and the stroke cycles respectively. Expressions for the net rates are calculated according to the principles in (40) (refer to supplemental info for a detailed derivation):

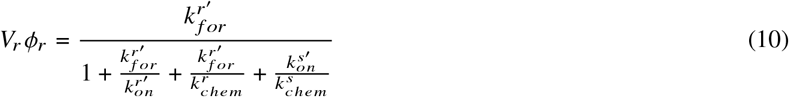

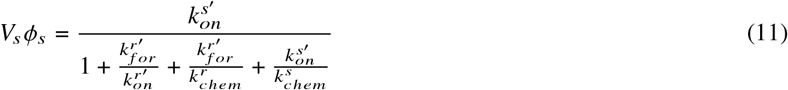

where

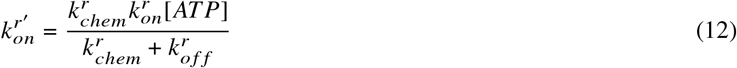

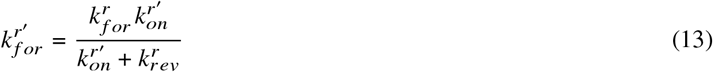

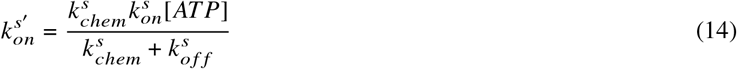

The solutions obtained from both the Gillespie simulations and the analytical methods agree consistently. Some examples are shown in Fig. S1.

### Exploring Rate Space

To sample a representative region of the phase space that a motor could access we varied the rate constants 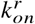 and 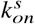 logarithmically in the ranges [0.1,10] pM^−1^s^−1^ and [0.1,100] pM^−1^s^−1^ respectively.

We varied 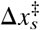 linearly in the range [0.01,1] nm. At every unique combination of rate constants and at every force we calculated the average velocity from the analytical solution. We then used the peak finding algorithm implemented in Scipy 1.2.1 (41) to find a peak in the average velocity as a function of force for every combination of the above mentioned parameters.

### Dwell Time Analysis

We obtained the dwell time distributions from the stochastic simulations of the motor by recording the waiting time between two consecutive steps taken by the motor, *τ*. We defined this as the time between two consecutive ATP hydrolysis steps. The randomness parameter was calculated from these distributions according to (42):

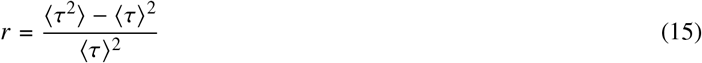

We repeated this for a range of forces to obtain the behavior of the randomness parameter as a function of force.

**Code Availability** Code for performing the Gillespie simulation, calculating average velocity for a range of forces and ATP concentrations, calculating the randomness parameter and visualizing the results can be found at https://wustl.box.com/v/2020-parallel-motor-mech

## RESULTS

### Construction of a parallel ratchet/stroke motor mechanism

To study the behavior of a hypothetical motor that has access to both ratchet and stroke mechanisms, we built a parallel path kinetic scheme (Fig. 1). The top path describes a ratchet mechanism where thermal fluctuations allow the motor (M) to explore positions N and N+1 according to the forward and reverse translocation rate constants 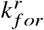 and 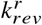. Here, ATP may bind to and dissociate from the post-translocated state (*M_n_*_+1_) with the the rate constants [ATP] 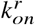 and 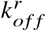 respectively. From the ATP-bound post-translocated state (*M_n_*_+1_ATP), ATP hydrolysis with rate 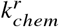 completes the cycle. The bottom path describes a stroke mechanism where ATP binding and dissociation occur in the pre-translocated state (*M_n_*) according to the rate constants 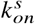 AND 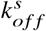 respectively. Subsequently, the cycle is completed via ATP hydrolysis and coupled forward translocation with a rate 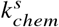.

### Average velocity and fractional flux

We first explored how the overall velocity of this motor depends on individual rate constants. In the case of a parallel mechanism, the motor velocity is determined by a flux-averaged net rate constant along the two paths for stepping. The net rate constant collapses all intermediate steps in a kinetic path to single kinetic rate constant describing the average stepping time along that path (40) (see Methods). Depending on the relative magnitudes of the net rate constants describing each individual path, the motor steps more frequently according to one or the other mechanism. To demonstrate this, we calculated the fractional flux through the ratchet arm (*ϕ*_*r*_) and average velocity of the mechanism (⟨*V*⟩) shown in the inset of Figure 2A for a range of net rate constants. The net rate constants of the ratchet path 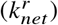 and stroke path 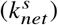 were varied by varying the rate constants 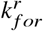, 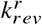 and 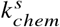 in an exponential range.

**Figure 2:**
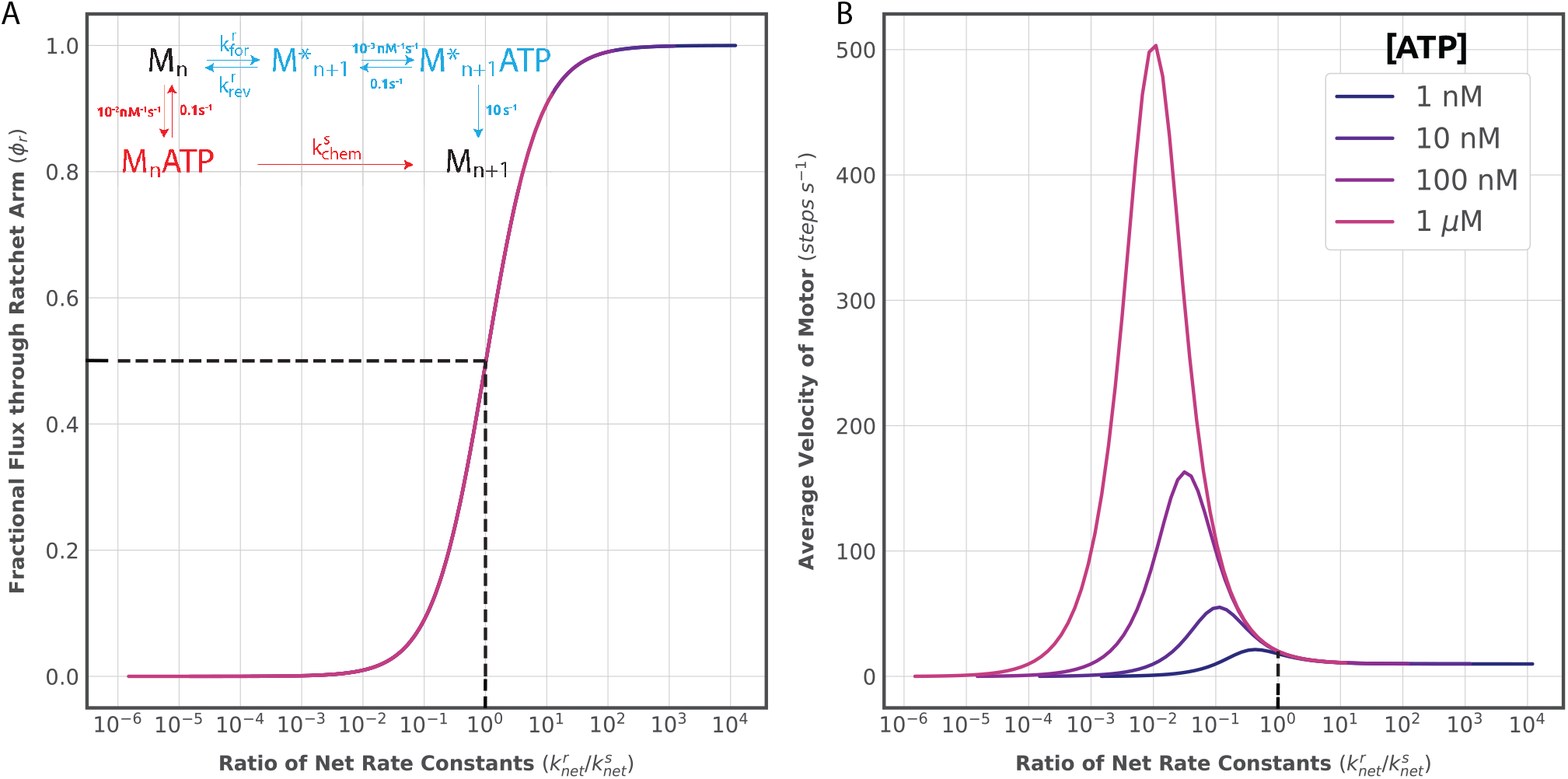
**(A)** Fractional flux through the ratchet arm of the parallel pathway in the presence of different ATP concentrations (see legend in 1B). The curves cannot be distinguished as they overlap exactly. **(B)** Average velocity of the motor as a function of the ratio of the net rates constants for varying [ATP]. Dashed lines on both panels indicate the point where fractional fluxes through both the ratchet and the stroke are exactly 0.5.

We further repeated this for a range of ATP concentrations. The dependency of the preference of the motor to step in either path on 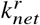 and 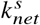 can most clearly be seen by plotting *ϕ*_*r*_ as a function of the ratio 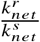 (Figure 2A). This curve shows the fractional flux through the ratchet arm of the pathway for varying concentrations of ATP in the system. We see that the fractional flux is not sensitive to the concentration of ATP for any given ratio of net rate constants but increases as magnitude of 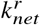 increases relative to 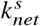.

By comparing the average velocity as a function of net rate constant ratio (Fig. 2B) and the fractional flux plot, it becomes clear that as the flux transitions from stroke to ratchet paths, a peak develops in the average velocity. Although here we are artificially manipulating the rate constants of the model, this interesting behaviour suggested to us that mechanochemical variables that affect the rate constants of each path differently may also lead to peaked velocity profiles.

### Fractional flux as a function of ATP concentration

Molecular motors require an energy source such as a nucleotide triphosphate molecule (often ATP) in order to perform work. Therefore, ATP concentration ([ATP]) changes motor behavior and is often used as a straightforward experimental variable to manipulate in order to test hypothetical mechanisms. Further, molecular motors are enzymes hydrolyzing their energy sources to drive conformational changes. In this regard, the parallel motor mechanism is analogous to a parallel ligand binding mechanism where ATP is the ligand. This must mean that [ATP] can change the fractional flux of the motor through either pathway in the context of a single set of rate constants as in (35). Indeed, we see this behavior (Fig. 3A (inset). We explored if this property could be used to experimentally distinguish a parallel mechanism from the behavior of either mechanism alone. Fig. 3A shows the average velocity as a function of [ATP] obtained from the analytical steady state solution of the system. Qualitatively, we see that the curves for both the parallel and individual mechanisms show a monotonic increase and saturation. In fact, for this combination of rate constants (see inset), the curve for the linear stroke mechanism overlaps with that for the parallel mechanism thus eliminating any possibility of distinguishing the two mechanisms using the [ATP] titration alone. To explore this more rigorously, we looked at the analytical expression for the average velocity of the motor in the parallel mechanism as a function of [ATP].

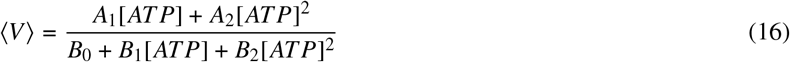

Where, *A*_1_, *A*_2_, *B*_0_, *B*_1_, *B*_2_ are various combinations of the individual rate constants of the reactions described in detail in equations 12-16 in the supplemental material.

**Figure 3:**
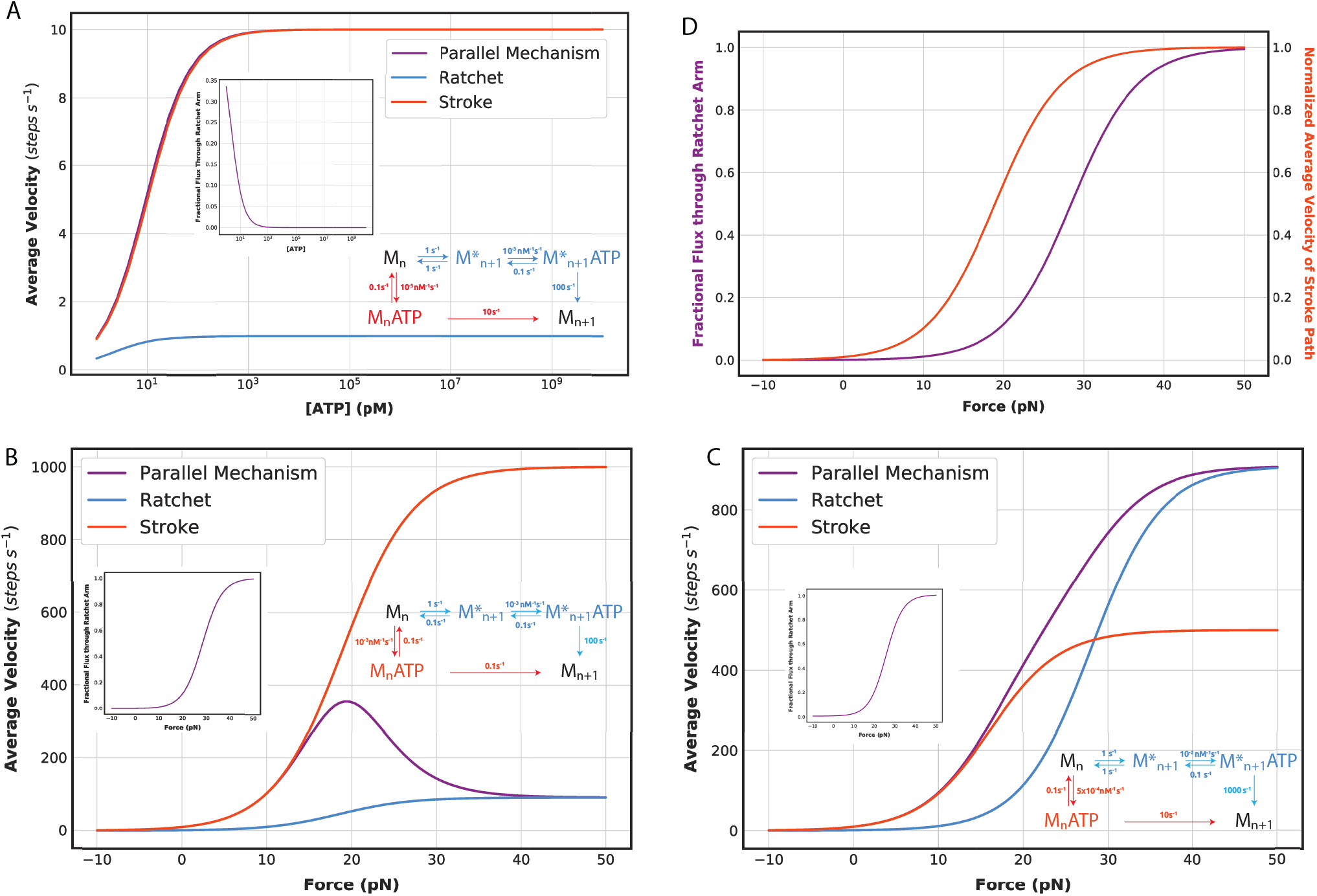
Peak Observed in Average Velocity as a Function of Applied Force. **(A)** Average velocity of the motor shown in the scheme in B as a function of [ATP] for each individual arm and the parallel mechanism. **(inset)** Fractional flux through the ratchet arm as a function of [ATP] **(B)** A peak is observed when the maximum velocity of the Ratchet path is slower than that of the Stroke path. [ATP] = 1 nM, 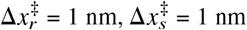, Δ*x* = 2 nm **(C)** No peak is observed when the maximum velocity of the ratchet path is faster than that of the stroke path even as the fractional flux shifts to the ratchet path at higher forces (inset). [ATP] = 1 nM, 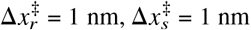, Δ*x* = 2 nm **(D)** Average velocity of the stroke path (normalized to 1) is plotted as a function of the applied force (purple) and compared against the fractional flux through the ratchet arm (red) for the mechanism shown in **B**

Equation (16) has a form similar to the Michaelis-Menten steady-state velocity equation for linear enzyme mechanisms (43) with additional second order [*ATP*] terms. At [*ATP*] < 1*M* these terms would also scale monotonically and would have a smaller contribution than the first order [*ATP*] terms. Therefore, one would not expect a qualitatively different curve for any combination of rate constants. Furthermore, distinguishing the mechanisms by fitting experimental data to these two forms can also be inconclusive as any curve that fits the Michaelis-Menten equation could fit better to the equation for the parallel mechanism due to the additional parameters. This means that [ATP] titrations, which are often used to elucidate the mechanism of motor action, would not generally be a reliable way to identify a parallel mechanism. This prompted us to look at force as a variable that might lead to a unique behavior of the parallel mechanism.

### An optimum force and a global maximum in the average velocity

All molecular motors are sensitive to force. Since distance along a track can be used as a reaction coordinate for these enzymes, force changes the relative free energy between states depending on the interstate distance. This results in force-dependent rate constants that describe changes in motor position. In a Brownian ratchet mechanism, force affects the thermally activated transitions between pre- and post-translocated states. In a power stroke mechanism, force affects the rate of the chemical step coupled to motion. In general, in the case of our parallel mechanism motor, each path will have a unique force-dependence. Since the probability of the motor stepping via one or the other paths depends on the relative magnitudes of the net rate constants for each path, we reasoned that in addition to modulating the overall velocity of the motor, force would modulate the probability of using a particular mechanism. That is to say that force should change the fractional flux just as ligand concentration does in parallel ligand binding mechanisms (35). Using a set of rate constants with pre-determined force dependencies (i.e. fixed transition state distances), this can be shown to be the case (Fig.3B (inset)). Specifically, opposing force favors the power stroke pathway while assisting force favors the Brownian ratchet pathway. This effect can be rationalized by the fact that greater assisting force (or lesser opposing force) increases the probability of ATP binding in the ratchet by promoting formation of the post-translocated state while not affecting the probability of ATP binding in the stroke.

In general, the direct observation of fractional fluxes in protein pathways is challenging. However, in the case of ligand binding, non-monotonic observed kinetics of equilibration can sometimes reveal a shift in fractional flux experimentally (35). To begin with, we asked whether some feature of the force-velocity curve might serve as an indicator of the shift in fractional flux. Surprisingly, for certain sets of rate constants, the velocity no longer depended monotonically on force as would be expected for an individual mechanism. Instead, the force-velocity relationship contained a global maximum (Fig. 3B). As increasing assisting force is applied to the system, the velocity begins to increase as would be expected. However, at larger assisting forces, the velocity decreases. By comparing the fractional flux and the force-velocity curves (Fig.3B (inset)), one can see that the peak in velocity occurs concomitantly with the transition in flux.

In the case of ligand binding, the experimental indicator of parallel paths (i.e. a non-monotonic observed rate) is only observable with a subset of possible combinations of rate constants even though the shift in fractional flux is always occurring (35). For this reason, we asked under what sets of rate constants the peak in motor velocity would be observable. In particular, the peak is a consequence of sequential transitions in the motor mechanism as a function of force. First, the saturated velocity of the ratchet path at high assisting forces must be lower than that the saturated velocity of the stroke path (Fig.3B and C). Only in this case will the average velocity decrease as the flux transitions to the ratchet path. Second, the force dependence of the flux must occur at larger forces as compared to the force dependence of the stroke path velocity (Fig.3D). This allows for the average velocity to increase appreciably prior to becoming dominated by the ratchet path. The converse case, where the flux increases rapidly before the average velocity of the stroke, leading to no observed peak in the velocity is shown in Fig. S2.

### Regions of parameter space that lead to peaked velocities

While these rules capture the qualitative determinants of whether a peak velocity exists, the analytical expression for the average velocity can be used to quantitatively explore the phase space for any set of rate constants. Using a peak finding algorithm (see Methods), we sparsely explored rate space and plotted the results according to whether a particular set contained a peak velocity (Fig. 4). The results are consistent with the qualitative expectations described above. At the lowest ATP concentration (Fig. 4A) As the stroke gets faster 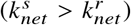 we observe a peak, provided stroke has a greater force sensitivity i.e., 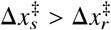. We note that it is also possible to have peaks located in the opposing force regime for particular combinations of rate constants (Fig. S3). This is because, fractional flux and average velocity are both continuous functions of force and the position of the *F* = 0 point in these curves is simply a reflection of the basal rate constants. In other words, these curves can be shifted along the force axis by changing a subset of the basal rate constants. Further, increasing [ATP] increases the possibility of a peak occurring as it increases the flux through the stroke path at low or zero opposing force.

**Figure 4:**
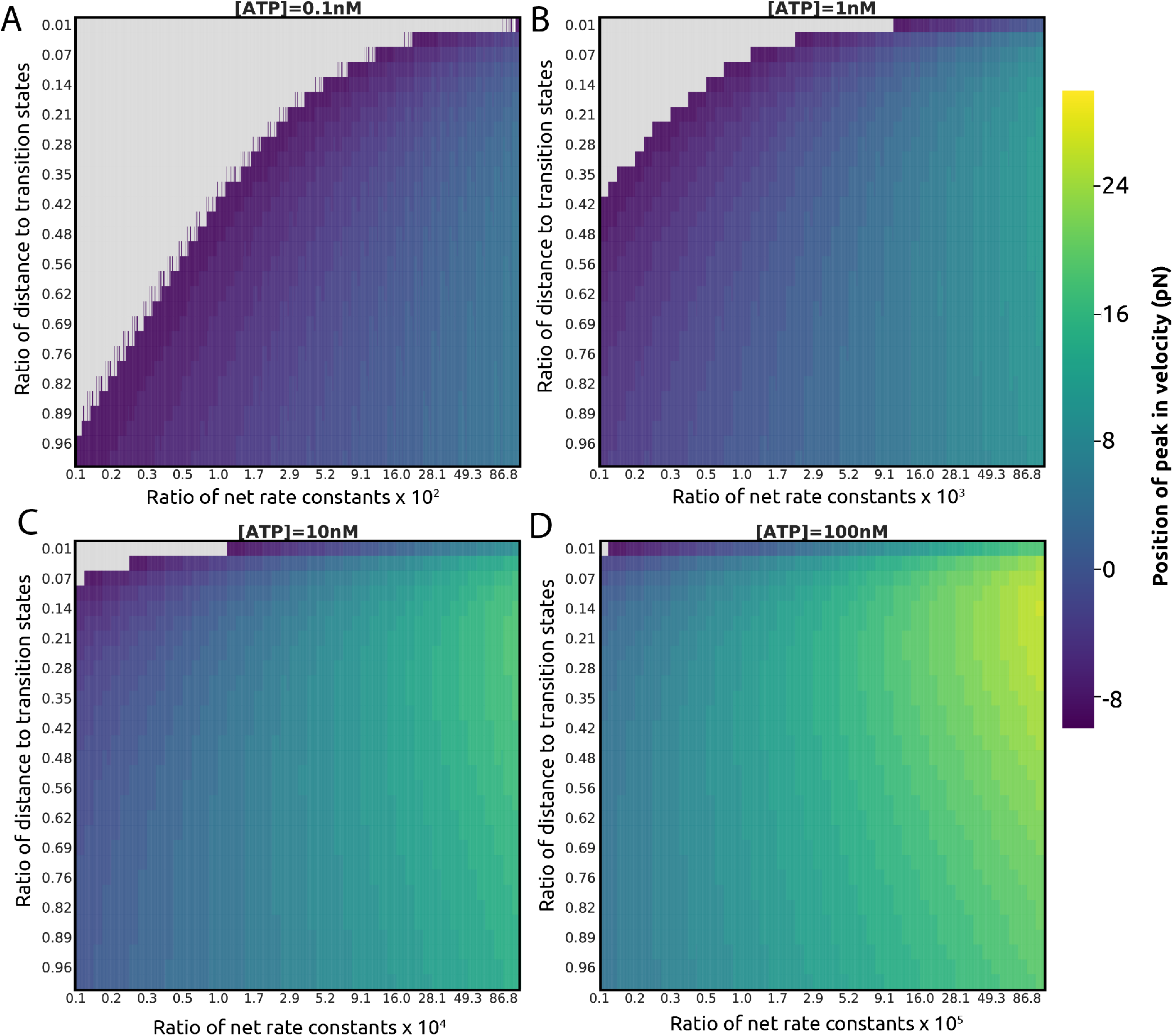
Exploring Phase Space Accessible with the Parallel Pathway. Force at which peak in average velocity occurs in pN for a given combination of rate constants (quantified by the ratio of the net rate constants of the individual stroke and ratchet paths at zero force, 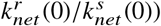 and for different force dependence of each path (quantified by ratio of the distance to transition states in each individual path, 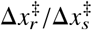) at [ATP] = 0.1 nM **(A)**, 1 nM **(B)**, 10 nM **(C)**, 1 μM **(D)**. Grey space indicates the region of the phase space where a peak is not observed. The mechanism is shown in Fig S4.

### The randomness parameter as an indicator of parallel pathways

We have shown that a force-dependent fractional flux would be detectable experimentally for a subset of rate sets via a peak in the average velocity. However, for other rate sets, the existence of parallel paths would go undetected as the force-velocity curve remains monotonic. Since molecular motors are frequently studied via single-molecule methods we asked if single molecule analyses would be better able to detect the presence of a parallel path mechanism. Specifically, analyses of dwell time distributions and their force-dependence are routinely used to uncover underlying mechanisms (2, 44, 45). More specifically, the randomness parameter, which is essentially the inverse of the variance of the dwell-time distribution, can be used to estimate the number of underlying rate-limiting steps (46). We performed stochastic simulations using the Gillespie algorithm to simulate single-molecule stepping data for the parallel path mechanism and populated dwell time histograms at a range of forces (Fig. 5A). For each individual path alone, the randomness parameter showed a single minima (Fig. 5B). We reason that this behavior is due to a change in the rate-limiting step of the pathway as a function of the force. As the assisting force increases, the force dependent step limits the rate as opposed to the enzymatic steps. The inverse of the randomness parameter is proportional to the number of rate-limiting steps for linear pathways (46). Therefore, a dip in randomness parameter would indicate a transition where there are a maximum number of rate-limiting steps. However, for the parallel mechanism, and in analogy to the non-monotonicity of the average velocity described above, a peak can be observed in the randomness parameter. In fact, this signal of the parallel mechanism is more sensitive than that of the average velocity as can be seen when the two observables are compared side-by-side for a given set of rates (Fig. 5C). Specifically, there are cases for which the velocity increases monotonically, while the randomness parameter shows a peak (Fig. 5D), thus increasing the range of rate constant sets for the parallel mechanism that would be detectable experimentally.

**Figure 5:**
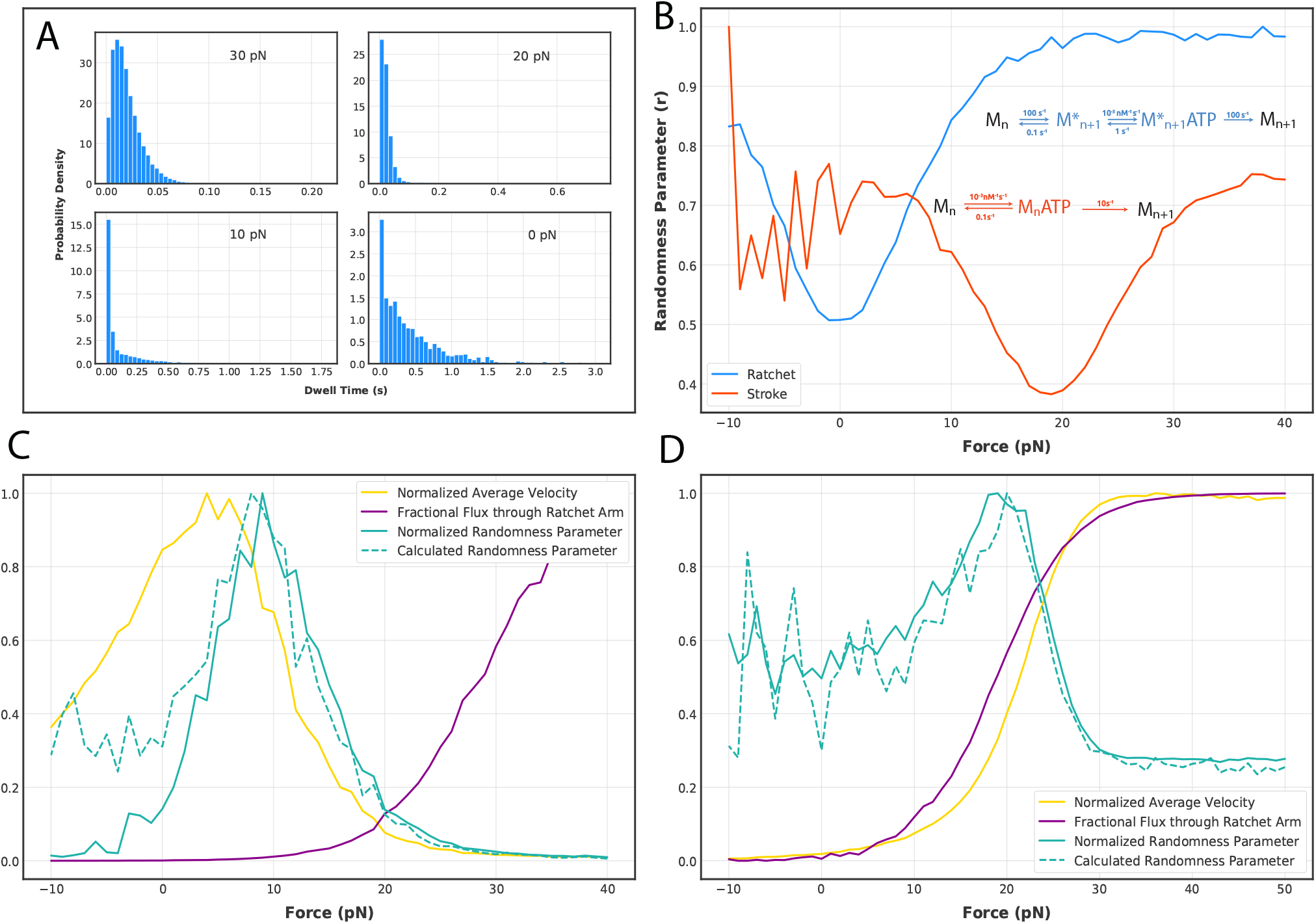
Randomness Parameter can be an Indicator of Parallel Pathways. **A)** Example of dwell time distributions obtained at different forces for a given set of rate constants and [ATP]. **B)** Randomness parameter (r) as a function of force for a linear ratchet and a linear stroke motor shown in blue and orange respectively. [ATP] = 0.1 nM (ratchet) and 10 nM (stroke), 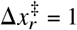 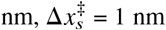, Δ*x* = 2 nm **C)** A peak is observed in r (green) as a function of force similar to the peak in average velocity (yellow). r and velocity are both normalized to 1. **D)** Example of a set of rate constants where a peak is observed in r whereas velocity shows a monotonic behavior. Mechanisms used in **C** and **D** are shown in Fig S4

We then explored if we could obtain an analytical expression for the randomness parameter similar to the ones derived for the velocity as a function of force and [ATP]. However, analytical solutions for the randomness parameter for non-linear schemes are highly non-trivial. We turned to an expression derived for the randomness parameter for a polymerase walking on a heterogeneous substrate such as DNA or RNA made of four different bases (42). The authors reasoned that the dwell time distributions obtained from a single molecule analysis of such a motor would be a weighted average of the distributions for the corresponding homogeneous substrates. Therefore, the randomness parameter would be given by

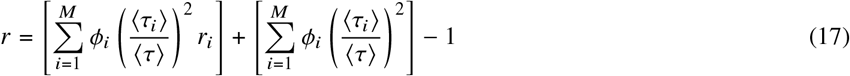

Where *ϕ*_*i*_ is the fractional flux, ⟨*τ*_*i*_⟩ the average dwell time and *r_i_* the randomness of the *i*th pathway and *M* is the total number of possible pathways in the mechanism.

The parallel mechanism we discuss in this work is analogous to the above in that the net dwell time distribution should be a fractional flux weighted average of the dwell time distributions for the linear ratchet and stroke mechanisms. We demonstrate the utility of this expression in Figs. 5C,D. Here, the dotted lines show the net randomness parameter for a range of forces for the parallel mechanisms calculated from the Gillespie simulations of the corresponding individual stroke and ratchet pathways (i.e. Fig. 5B). We see good overlap of the calculated and simulated randomness parameters. Therefore, we reason that an experimentally obtained r vs. force curve could be fit to equation 17 to obtain the flux and randomness of the individual pathways. The flux thus obtained can also be used to constrain the velocity analysis such that the average velocities of the individual pathways can be obtained with more confidence even when no peak is observed in the velocity vs. force curves. However, as is the case for ligand-binding, there are still many sets of rate constants for the parallel mechanism that do not result in a detectable signal in either the average velocity or the randomness parameter (Fig. S5).

### The lack of experimental evidence for parallel paths cannot be used to exclude their existence

For a large region of rate space, the behavior of the parallel path mechanism does not provide any feature that would serve to identify its presence. In particular, for these rate sets, both the force-velocity curve and the dwell-time distributions can be well-fit by linear mechanisms. This is consistent with expectations that more complex models can always fit data that are well described by simpler models. Especially in this case as the parallel mechanism is actually constructed from the simpler linear models typically used to describe motor function. However, we caution that the simplest model that can account for the data is not necessarily the best model to describe the actual underlying function of a molecule.

## DISCUSSION

We have demonstrated that, for a molecular motor with access to either Brownian ratchet or power stroke paths ATP concentration and external bias can modulate the fractional flux through each mechanism. Furthermore, we report that for a subset of possible rate constants, the shift in fractional flux through either linear arm caused by force may lead to an optimum velocity of the motor. In contrast, the average velocity as a function of ATP concentration remain monotonic. We also show that the randomness parameter, obtained from the distribution of single-molecule dwell-times, shows a peak as a function of force for a broader range of rate constants compared to the average velocity. This peak could be used to directly identify a parallel mechanism. However, we note that there still remains a large region of the rate space where neither the average velocity nor the randomness parameter would show an unambiguous signature of a parallel mechanism.

Whether molecular motors exist which are capable of functioning according to the model proposed here is unclear. In our survey of the literature, we did not encounter examples of non-monotonic force-velocity or force-randomness parameter curves. However, since there are many rate constant combinations which would not lead to these observations, the question may still be a valid one. Additionally, it may be that these curves have not been explored with sufficient force resolution or precision to determine the presence of non-monoticity in a statistically significant manner.

In conclusion, we note that the folding, ligand-binding, catalysis, and movement of macromolecules are determined by a many-dimensional energy landscape. In principle then, they will have a multitude of possible paths connecting any two states on that landscape. In this spirit, we have explored the hypothetical behavior of a molecular motor that can function as either a power stroke or a Brownian ratchet and have presented ways that this might be observable experimentally. We have also pointed out that this underlying mechanism may exist without a clear way to distinguish it from a simpler linear mechanism. Therefore, we suggest that, just as conformational selection and induced fit may coexist, the ratchet/stroke duality may be a false one. In particular, in some cases, the answer to the question of whether a molecular motor uses a Brownian ratchet or a power stroke mechanism may simply be, “both”.

## AUTHOR CONTRIBUTIONS

E.A. Galburt designed the research. U.L. Mallimadugula carried out all simulations and analyzed the data. E.A. Galburt and U.L. Mallimadugula wrote the article.

## ACKNOWLEDGMENTS

We acknowledge the support of the Biochemistry and Molecular Biophysics department (EAG) and the Biochemistry, Biophysics, and Structural Biology program within the Division of Biology and Biomedical Sciences graduate program (ULM), both at the Washington University School of Medicine.

